# PartitionFinder-mAIC: Phylogenetic Partitioning using Marginal Akaike Information Criterion

**DOI:** 10.64898/2026.09.04.749328

**Authors:** Huaiyan Ren, Thomas K.F. Wong, Changsen Jiang, Edward Susko, Robert Lanfear, Bui Quang Minh

**Affiliations:** School of Computing, College of Systems & Society, Australian National University, Canberra, ACT 2600, Australia; Mathematical Sciences Institute, College of Systems & Society, Australian National University, Canberra, ACT 2600, Australia; Ecology and Evolution, Research School of Biology, College of Science & Medicine, Australian National University, Canberra, ACT 2600, Australia; Department of Mathematics and Statistics, Faculty of Science, Dalhousie University, NS, Canada

**Keywords:** phylogenetics, partition model, model selection, partitioning scheme, marginal Akaike information criterion, marginal likelihood, IQ-TREE

## Abstract

Partition models are widely used in phylogenomic analyses to account for heterogeneous evolutionary processes across different regions or loci of a sequence alignment. How alignment regions are grouped into partitions (the partitioning scheme) affects both the degree of over- or under-parameterization and the accuracy of downstream phylogenetic inferences. PartitionFinder is a widely adopted framework for selecting an optimal partitioning scheme. Using Akaike information criterion (AIC) and Bayesian information criterion (BIC), PartitionFinder merges partitions with similar evolutionary processes to avoid model overfitting. However, AIC and BIC are based on conditional likelihoods that treat partition assignments as fixed, whereas the recently introduced marginal AIC (mAIC) averages site likelihoods over the models of all partitions, providing a more appropriate criterion for inferring global parameters such as tree topology and branch lengths (Susko et al. 2026). Here, we implement mAIC for partition models in IQ-TREE 3 and integrate it into the PartitionFinder algorithms. Using a range of simulated and empirical DNA and protein datasets, we show that PartitionFinder-mAIC yields partitioning schemes with fewer partitions than those selected by AIC or BIC, and additionally improves phylogenetic inference at most key branches of the green plant evolution. The new PartitionFinder-mAIC is available in IQ-TREE version 3.1.4 with the command-line option -merit mAIC.

## Introduction

In phylogenomic analyses of multi-locus or genome-scale datasets, sequence alignments are highly heterogeneous. Different sites, genes, loci, or protein domains can evolve under distinct substitution processes, rates, and compositions of nucleotides or amino acids (Lartillot and Philippe 2004; Nylander et al. 2004; Misof et al. 2014). Accounting for this heterogeneity is important as failure to do so can lead to biased tree topology, branch length, and branch support (Anderson and Swofford 2004; Brown and Lemmon 2007; Kainer and Lanfear 2015).

Partition models are commonly used to capture across-site heterogeneity in likelihood-based phylogenetics. A partition model divides an alignment *a priori* into subsets (e.g., by gene, codon position, exon, intron, or protein domain) and assigns an independent substitution model to each subset (Nylander et al. 2004; Lanfear et al. 2012). A practical difficulty with partition models is choosing an appropriate partitioning scheme. Common user-defined schemes based on gene boundaries or individual codon positions within individual loci are often too fine-grained (Ren et al. 2025). Assigning a separate model to each of hundreds or thousands of loci usually leads to overparameterization and can reduce phylogenetic accuracy (Brandley et al. 2005; Lanfear et al. 2012; Susko and Roger 2020). PartitionFinder (Lanfear et al. 2012; Lanfear et al. 2017) addressed this problem by iteratively merging subsets with similar evolutionary processes. Here, pairs of partitions are merged if this improves the model fit, typically measured by Akaike (Akaike 1974) or Bayesian (Schwarz 1978) information criterion (AIC or BIC). PartitionFinder has been implemented in IQ-TREE (Chernomor et al. 2016; Wong et al. 2026) and is now widely adopted in phylogenomic studies (Cannon et al. 2016; Stiller et al. 2024).

The information criteria used by PartitionFinder are computed from the likelihood of the partition model. In a partition model, each site is assigned to a fixed partition, and its likelihood is evaluated conditionally on the substitution model, tree topology and branch lengths of that partition, giving a conditional likelihood. However, when the quantity of interest is a global parameter such as the tree topology or branch lengths, the appropriate likelihood is the marginal likelihood, computed by averaging each site’s likelihood over the models from all partitions rather than conditioning on a single fixed assignment (Susko et al. 2026). This is because the partition assignment of each site is itself uncertain, and conditioning on a fixed assignment ignores this uncertainty, which can result in poor topological and global parameter estimation even in cases with large amounts of data (Susko et al. 2026). The marginal AIC (mAIC; Susko et al. 2026) is based on marginal likelihoods and provides the conceptually appropriate criterion for model selection when global phylogenetic parameters are the target of inference. This has been validated on empirical amino acid datasets (Ren et al. 2026). However, mAIC is calculated once the partitioning scheme was already determined by AIC or BIC. This raises the question of whether a better fit partitioning scheme can be found by directly applying mAIC or not. Currently there is no such tool available to answer this question. In the following, we refer to AIC computed on conditional likelihoods as cAIC to distinguish from mAIC.

To address this gap, we here extend PartitionFinder to include mAIC as the criterion for partition merging. This is not as simple as it may sound. Because cAIC evaluates each site only under the model of its assigned partition, it does not require information from other partitions. Computing mAIC, by contrast, requires re-evaluating each site’s likelihoods under the models from all other partitions in the scheme. Replacing cAIC with mAIC in the PartitionFinder merging algorithm is therefore non-trivial: at each merging step, the mAIC of every candidate merged scheme must be recomputed from scratch, naively adding substantial computational overhead to an already expensive model search.

Here we solve this problem and integrate mAIC into the PartitionFinder algorithm in IQ-TREE, called PartitionFinder-mAIC. We describe how the relaxed clustering (Lanfear et al. 2014) and greedy search strategies (Lanfear et al. 2012) can be modified to use mAIC as the merging criterion with only modest additional computation. Using simulated and empirical DNA and protein datasets, we then compare the partitioning schemes and resulting phylogenetic trees obtained under cAIC, mAIC, and BIC. We show that mAIC-based PartitionFinder consistently yields partitioning schemes with fewer partitions than cAIC or BIC, a result supported by both simulated and empirical datasets. We demonstrate that in simulated datasets the choice of optimisation criterion (cAIC, BIC, or mAIC) has little effect on inferred tree topologies. However, on empirical datasets models selected with the PartitionFinder-mAIC yielded phylogenies with greater topological concordance with established references and stronger bootstrap support than those obtained under cAIC or BIC. The new feature is available in IQ-TREE version 3.1.4 (Wong et al. 2026).

## Materials and Methods

### Implementation of mAIC computation for partition models

We implemented mAIC computation as described in Susko et al. (2026). Given a concatenated alignment *D* with a total of *N* sites in *K* partitions *D*_1_, … , *D_K_* (*D_i_* is the set of sites in partition *i* ), a partition model Θ is a set of trees *T* = {*T*_1_, … , *T_K_*} and models (substitution model rate matrices, state frequencies and rate heterogeneity across sites) *M* = {*M*_1_, … , *M_K_*}. The conditional likelihood of the partition model is:

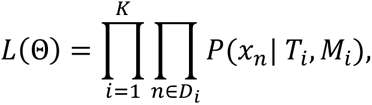

where P(*x_n_*|*T_i_*, *M_i_*) is the probability of site pattern *x_n_* in partition *D_i_* , conditional on it evolving under *T_i_* and *M_i_*. Here, *x_n_* = {*x*_1*n*_, *x*_2*n*_, … }, where *x_sn_* is the character state for the s^th^ taxon. If the partition model has df (degree of freedom) as the number of free parameters, the cAIC and BIC of partition models are calculated as *cAIC* = −2ln*L*(Θ) + 2df and *BIC* = −2ln*L*(Θ) + df × ln*N*.

To compute the marginal likelihood, for each site pattern *x_n_* in partition *D_i_* , we need to compute its likelihood if it evolved under *T_j_* and *M_j_* of other partitions *j* (*j* ≠ *i*): *P*(*x_n_*| *T_j_*, *M_j_*).

The marginal probability of *x_n_* is computed as the weighted sum over all these probabilities, where the weight *w_j_* is the proportion of sites in *D_j_* over all sites *N* in the concatenated alignment *D*. Therefore, the marginal likelihood of the partition model is:

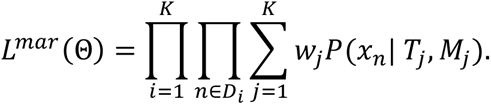

Here, the double product over partitions and sites 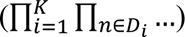 is equivalent to a single product over all *N* sites 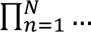 in the alignment. In our implementation, we retain the partition-wise double-product form, as it allows the marginal site likelihoods to be computed independently for each partition and thus can be parallelised across threads, improving computational efficiency. The mAIC is then computed as *mAIC* = −2ln*L^mar^*(Θ) + 2df.

Computing a site likelihood *P*(*x_n_* | *T_j_*, *M_j_*) for a site *x_n_* in partition *D_i_* under a tree *T_j_* and a model *M_j_* of a different partition *D_j_* raises a practical issue when the taxon sets *S_i_* and *S_j_* of the two partitions differ due to missing data. Taxa present only in *S_j_* do not affect the likelihood. If there are taxa that are present in *S_i_* but missing from *S_j_*, we treat their states as marginalised under the stationary distribution of *M_j_*, so that

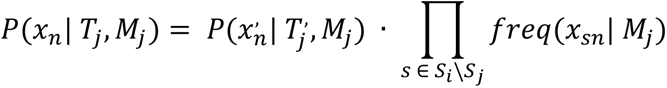

where *x_n_*, is the restriction of site *x_n_* to the taxa shared between the two partitions (*S_i_* ∩ *S_j_*), *T*,*_j_* is the subtree of *T_j_* restricted to the shared taxa. This treatment makes the mAIC computation compatible with partitioned alignments in which some taxa are missing from some partitions, which is common in phylogenomic datasets.

The mAIC computation in IQ-TREE supports parallelization across multiple threads. If more than one thread is assigned to the job, the likelihood computations for the partition pairs (*i*, *j*) will be automatically distributed across the available threads to accelerate the total computation.

The implementation is compatible with several partitioned-model settings in IQ-TREE: edge-linked (edge-equal and edge-proportional) and edge-unlinked partition models, reversible and non-reversible substitution models, and partitioned alignments containing multiple data types (DNA, protein, codon, etc.). For partitioned alignments that contain multiple data types, the marginal log-likelihood is computed separately for partitions that share the same data type and then summed across data types; the mAIC is computed from this total marginal log-likelihood.

### Algorithm design of PartitionFinder-mAIC

The new PartitionFinder-mAIC in IQ-TREE builds upon two partitioning search strategies: a relaxed clustering algorithm (the default; Lanfear et al. 2014) and a greedy algorithm (Lanfear et al. 2012). Both algorithms iteratively evaluate candidate pairs of subsets for merging and accept a merge when the information criterion score improves. We extend both strategies to support mAIC as the merging criterion.

We summarize our algorithm below. It follows the general framework of the strict hierarchical clustering algorithm described in the early version of PartitionFinder (Lanfear et al. 2014). However, because our implementation builds upon the IQ-TREE version of PartitionFinder, which integrates ModelFinder (Kalyaanamoorthy et al. 2017) and other enhancements, certain algorithmic details differ from the original description. Our algorithm proceeds as follows:

1. Estimate an initial phylogenetic tree topology from an input sequence alignment using maximum parsimony criterion (MP; Fitch 1971) and nearest neighbour interchange (NNI; Robinson 1971) ;
2. Initialize a partitioning scheme in which all user-defined data blocks are assigned to separate subsets;
3. For each subset in the current partitioning scheme, estimate the substitution model (GTR+I+G model for DNA and LG+I+G model for protein by default) and partition rate, then compute the likelihood and cAIC score for that subset;
4. Compute the mAIC of the current partitioning scheme;
5. Generate all candidate pairs of subsets;
6. If the relaxed clustering algorithm is applied: calculate the distances (difference in log-transformed tree length by default) between all candidate pairs and retain those with the smallest distance, up to the top 10% of all pairs (default; set via --rcluster options) but no more than 10 × the number of user-defined blocks (default);
7. For each retained candidate pair, estimate the substitution model (GTR+I+G model for DNA and LG+I+G model for protein by default) and partition rate of the merged subset, then compute its likelihood and cAIC score;
8. Sort the retained candidate pairs in increasing order of ΔcAIC, defined as the difference between the cAIC of the merged pair (*i*, *j*) and the sum of cAIC scores of the two independent subsets *i* and *j*;
9. Traverse the sorted candidate pairs:

a. If the relaxed clustering algorithm is applied: compute the mAIC of the partitioning scheme resulting from merging *i* and *j* . If this mAIC improves on the current scheme, merge the pair and continue to the next candidate; otherwise, skip the pair. Any pair in which subset *i* or *j* has already been merged earlier in the same traversal is also skipped;
b. If the greedy algorithm is applied: compute the mAIC of the partitioning scheme resulting from merging *i* and *j*. After walking through all candidate pairs, merge the single pair that yields the greatest mAIC improvement;
10. Return to step 4 and repeat until no pairs are merged in step 9;
11. Select and estimate the substitution model and partition rate (running the full ModelFinder procedure) for each subset in the final partitioning scheme and compute information criteria scores.

Throughout the algorithm, the same site likelihood *L*(*x_n_* | *T_b_*, *M_b_*), where *n* ∈ *D_a_*, is required many times, because the partition pair (*a*, *b*) whose likelihood is being computed is in general distinct from the candidate merge pair ( *i* , *j*) being evaluated at a given step. To avoid redundant computation, we cache *L*(*x_n_* | *T_b_*, *M_b_*), where *n* ∈ *D_a_*, for all partition pairs (*a*, *b*) and reuse the cached values whenever the same pair is encountered again.

### Evaluating correlations between mAIC improvement and other metrics

In the algorithm design, we use log-transformed tree length (step 6) to sort and select the candidate pairs during merging. Lanfear et al. (2014) examined four metrics for partition merging: partition rate (equivalent to log-transformed tree length), substitution model rates, state frequencies, and the Gamma alpha parameter, and found none correlated clearly with merging improvements in cAIC or BIC. They nonetheless sorted candidate pairs by partition rate similarity, retaining the top 10% (relaxed clustering only) as default, a setting inherited by the IQ-TREE implementation.

Here, we assessed whether ΔmAIC (the mAIC improvement from merging a pair) is associated with each of these four metrics, and with ΔcAIC (step 8), which would indicate that a cheaper metric could pre-select candidate pairs. For each candidate pair we measured the log-transformed tree length distance, the substitution model rate and state frequency differences (Euclidean distances), the rate heterogeneity difference (absolute Gamma alpha difference), and ΔcAIC.

To generate these data, we randomly drew two non-overlapping subsets of 40 loci (39 for the green plant plastid protein dataset) from each dataset in Table 1, then ran the greedy algorithm merging partitions from 40 down to 1, recording ΔmAIC and all metric values for every candidate pair at each iteration. We quantified associations using the absolute Spearman’s rank correlation (ρ), chosen because the merging decision depends only on the ranking of pairs, not on the actual scores of the pairs. Joint distributions are visualised as scatter plots in Supplementary Figure S1.

**Table 1.** Empirical datasets used to benchmark PartitionFinder-mAIC.

| Dataset | Data type | Sites | Loci | Taxa | Reference |
| --- | --- | --- | --- | --- | --- |
| Metazoan | DNA (no 3<sup>rd</sup> codon) | 89792 | 424 | 78 | Cannon et al. 2016 |
|  | Protein | 44896 | 212 | 78 |  |
| Birds | DNA (no 3<sup>rd</sup> codon) | 189220 | 300 | 363 | Stiller et al. 2024 |
|  | Protein | 94610 | 150 | 363 |  |
| Green plants<br>plastid | DNA | 58347 | 234 | 360 | Ruhfel et al. 2014 |
|  | DNA (no 3<sup>rd</sup> codon) | 38898 | 156 | 360 |  |
|  | Protein | 19449 | 78 | 360 |  |
| Archaea | Protein | 17191 | 84 | 364 | Dombrowski et al. 2020 |

In this analysis, the modified IQ-TREE version with the early-stopping condition disabled, which allows the greedy algorithm to merge partitions all the way down to a single partition and records the metrics of each partition pair, is available at Data, Code and Software Availability.

### Testing PartitionFinder-mAIC with Simulation

We simulated alignments along a random 200-taxon Yule-Harding tree using AliSim (Ly-Trong et al. 2022). The tree was rescaled to three levels of average branch length (0.02, 0.004, and 0.001) and seven ratios of external to internal branch length (1, 2, 4, 8, 16, 32, and 64), spanning a range of evolutionary divergences and tree-inference difficulties. Each simulated alignment comprised either 8 partitions (for DNA) or 4 partitions (for protein, which are generally shorter), with 1000 sites per partition. Partition rates were drawn from a uniform distribution between 1 and 10 and then rescaled to a mean of 1, so that the fastest partition evolved ten times faster than the slowest. Each setting was replicated 30 times.

To mimic the over-partitioned starting point, each simulated partition was randomly split into 5 artificial subsets of 200 sites each, yielding 40 (DNA) or 20 (protein) input partitions for PartitionFinder. We ran PartitionFinder with both relaxed clustering (default) or greedy (-- merge greedy) algorithms under cAIC, BIC, and mAIC, as well as under the true and the no-merge schemes, and inferred a tree for each alignment with 1000 ultrafast bootstrap replicates (UFBoot; Hoang et al. 2018). The IQ-TREE command line for PartitionFinder is:

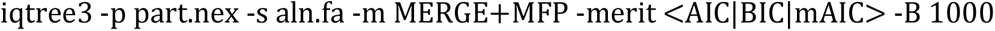

The command line for true or no-merge schemes is:

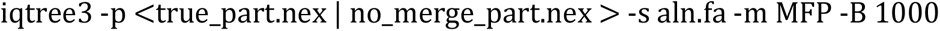

For each merging criterion, we recorded the number of partitions retained and evaluated tree inference against the true tree using the Robinson-Foulds (RF) distance (Robinson and Foulds 1981), the Lin-Rajan-Moret (LRM) distance (Lin et al. 2012), the Kuhner-Felsenstein (KF) distance (Kuhner and Felsenstein 1994), the difference in total tree length from the true tree, and the average UFBoot support of correctly and incorrectly inferred branches. Each criterion was compared against mAIC using paired Wilcoxon signed-rank tests (Wilcoxon 1945). Because all conditions were applied to the same simulated alignments, observations were paired within each condition combination. Tests were performed separately within each combination of mean branch length and branch-length ratio. To account for multiple comparisons across all cells and contrasts, p-values were adjusted using the Holm method (Holm 1979).

### Empirical datasets

We analysed eight empirical phylogenomic datasets spanning four taxonomic groups and multiple data types (Table 1): metazoan (Cannon et al. 2016), bird (150 loci subsampled from 14792 exon coding sequences; Stiller et al. 2024), green plant plastid (Ruhfel et al. 2014), and archaea (Dombrowski et al. 2020). DNA datasets were analysed with third codon positions excluded, except for the green plant plastid dataset, which was additionally analysed with all codon positions included. All datasets were also analysed as protein alignments where applicable.

### Applying PartitionFinder on empirical data

For each dataset, we ran the PartitionFinder relaxed clustering strategy using cAIC, BIC and mAIC. All analyses used the edge-proportional partition model, which is generally recommended over the edge-equal and edge-unlinked models (Duchêne et al. 2020; Ren et al. 2026). The IQ-TREE command line is:

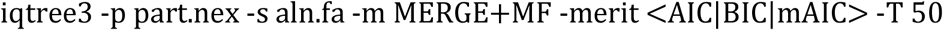

Each run was allocated 50 CPU threads. IQ-TREE applied efficient partition-level parallelization by assigning multiple threads to each partition proportional to its size. We recorded the runtime of each PartitionFinder step from the IQ-TREE log file, including model estimation during merging, final model estimation after merging, and mAIC score computation. We also extracted the memory estimates for initial model estimation, final optimization, and mAIC cache as reported by IQ-TREE. Total wall-clock time and peak resident memory were measured using the Linux /usr/bin/time utility.

To examine the behaviour of each criterion along the full merging path, we additionally ran the PartitionFinder greedy algorithm on a 40-locus metazoan DNA subset, merging partitions from 40 to a single partition (similar to the evaluating correlation section above) and recording cAIC, BIC, and mAIC scores at each step. The IQ-TREE command line is:

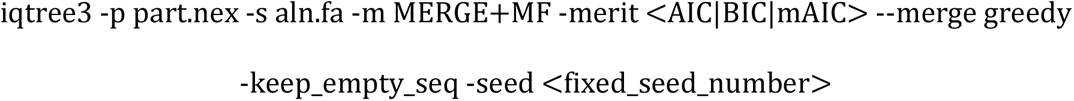

### Tree inference and comparison

We inferred Maximum Likelihood (ML) phylogenetic trees under each final partitioning scheme estimated by previous step and the original no-merge schemes on the green plant plastid datasets and applied 1000 ultrafast bootstrap replicates (Hoang et al. 2018). We compared the trees with published reference topologies. The command line for tree inference: iqtree3 -p <previous_step.best_scheme.nex|original_scheme.nex> -s aln.fa -B 1000

## Results

### No single metric is a strong proxy for mAIC improvement

No single metric showed a meaningful correlation with ΔmAIC: when candidate pairs were ranked by each metric across all iterations, replicates, and datasets, absolute Spearman’s ρ values were below 0.1 in every case (Supplementary Fig. S1). This was true not only for the four metrics examined by Lanfear et al. (2014) but also for ΔcAIC, indicating that none provides a reliable shortcut for ranking candidate pairs by mAIC improvement. For protein datasets, a single LG substitution rate matrix was applied to all partitions during the merging process, resulting in zero substitution rate distance for all partition pairs. This metric was therefore excluded from the protein analysis.

As all metrics showed no or weak correlation with the ΔmAIC throughout the merging process, we retained the default settings described in the Materials and Methods section for all subsequent analyses: candidate pairs are sorted and filtered to the top 10% by log-transformed tree length distance for the relaxed clustering algorithm (step 6) and by ΔcAIC for mAIC evaluation order (step 8), as ΔcAIC exhibited the highest correlation with ΔmAIC among all metrics tested.

### mAIC yields partitioning schemes with fewer partitions in simulated data

Figure 1 shows the number of partitions retained by PartitionFinder under each criterion across the simulated conditions, spanning different average branch lengths and ratios of external to internal branches. For DNA alignments, mAIC consistently retained fewer partitions than cAIC and BIC (Fig. 1A). For protein alignments, mAIC again retained fewer partitions than cAIC, and a number similar to BIC, though this varied across branch-length settings (Fig. 1B). BIC and mAIC retained close to the true number of partitions (8 for DNA and 4 for protein) when the mean branch length is 0.02 but retained fewer than the truth when branches were extremely short (≤0.004), whereas cAIC consistently merged fewer partitions.

**Fig. 1.**
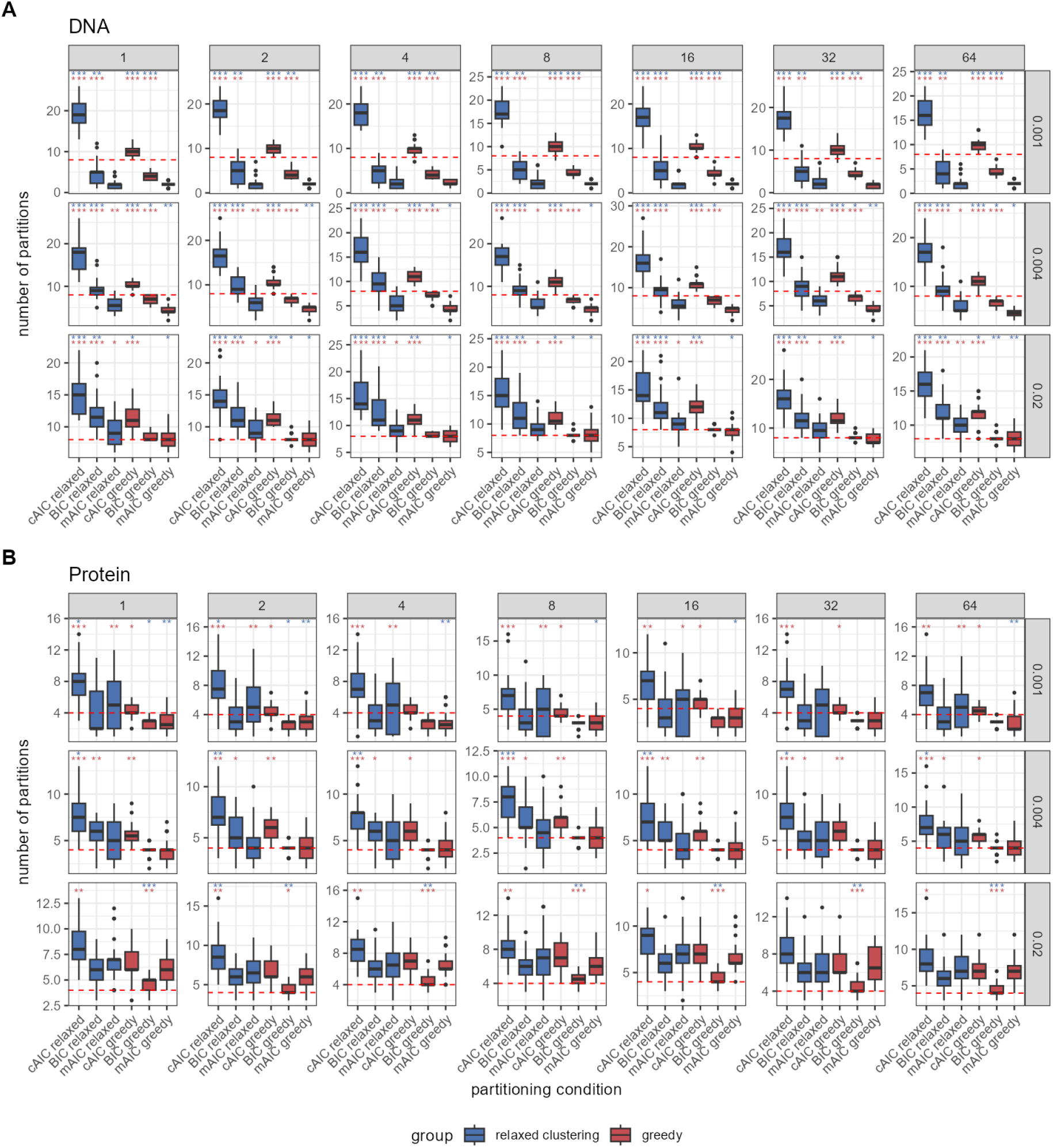
Number of partitions selected by each criterion and partition-merging strategy under simulation. (**A**) DNA and (**B**) protein datasets. Within each panel, boxplots show the number of partitions inferred across replicates, faceted by mean branch length (rows) and external-to-internal branch-length ratio (columns). Red dashed lines indicate the true number of partitions. Asterisks denote significant differences from paired Wilcoxon signed-rank tests, with each criterion compared against mAIC (blue, relaxed clustering; red, greedy). P-values were adjusted for multiple comparisons using the Holm method (∗ P < 0.05, ∗∗ P < 0.01, ∗∗∗ P < 0.001).

Across all conditions, the greedy algorithm retained fewer partitions than the relaxed clustering algorithm.

We then compared tree inference under each partitioning scheme against the simulated tree. Across all metrics (see Materials and Methods), mAIC did not differ significantly from the other two criteria (Supplementary Figs. S2–S7), with one exception: the difference in total tree length from the truth (paired Wilcoxon signed-rank tests). Trees inferred under mAIC schemes had smaller total tree length differences from the truth than those under cAIC or BIC. At average branch lengths of 0.02 and 0.004, this brought mAIC-based tree lengths closer to the true value, whereas at 0.001 the difference became negative, indicating that mAIC slightly underestimated the true tree length.

### mAIC yields partitioning schemes with fewer partitions in empirical datasets

Across all tested empirical datasets, mAIC-based merging produced partitioning schemes with fewer partitions than cAIC (Fig. 2A). For DNA data, cAIC retained the largest number of partitions (32-85% of the number of starting partitions), the BIC always retained fewer than the cAIC (12-38%) and the mAIC retained the fewest (11-22%).

**Fig. 2.**
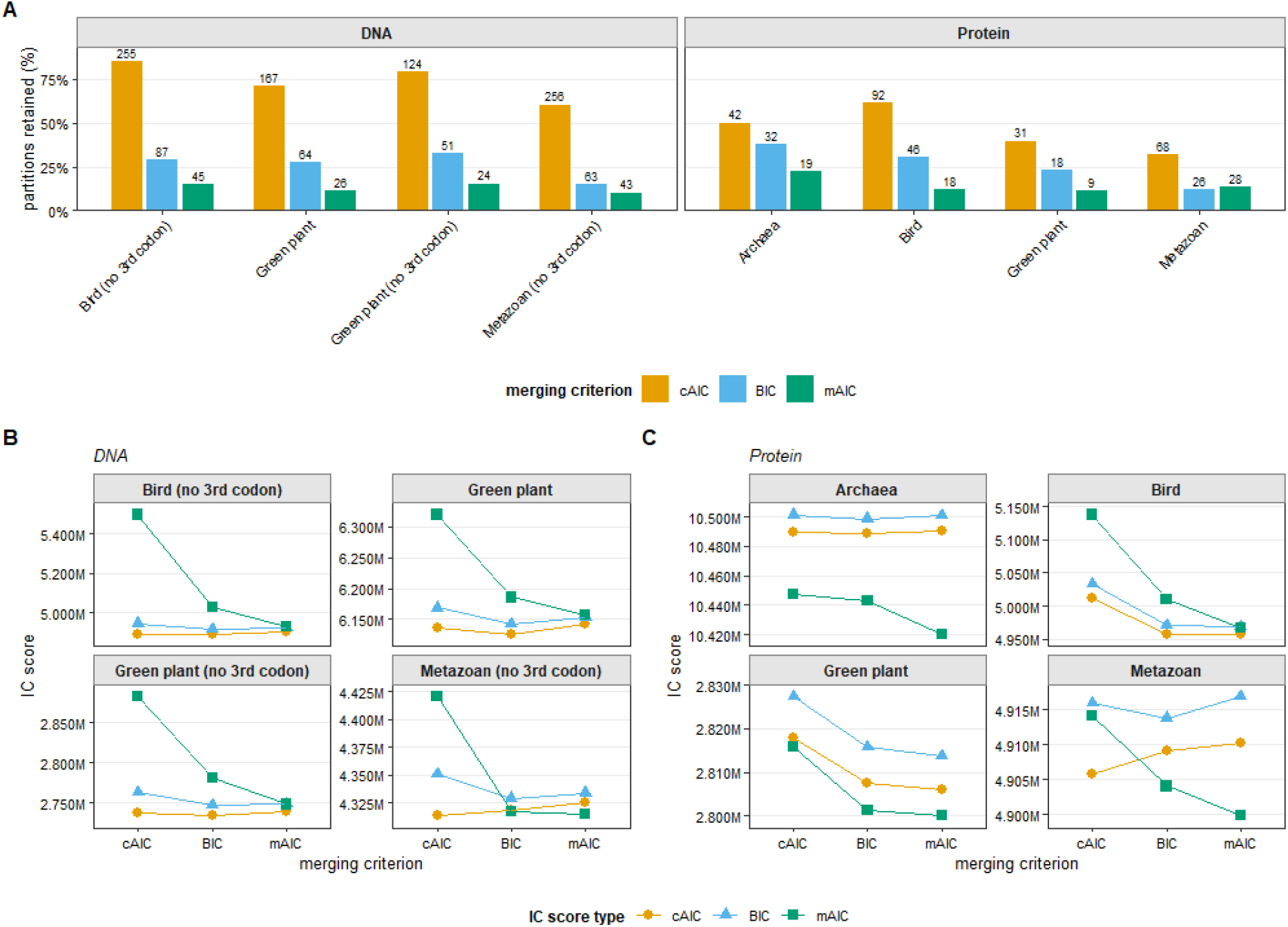
Partition model selection results under different merging criteria. **A**: Number of partitions retained by the PartitionFinder relaxed clustering algorithm. Numbers above each bar indicate the partition count. **B, C**: Information criterion scores of partition models estimated under the PartitionFinder relaxed clustering algorithm with different merging criteria for DNA (**B**) and protein (**C**) datasets.

For each partitioning scheme selected under a given merging criterion, we computed all three information criterion scores (cAIC, BIC, and mAIC). We compare schemes using the same information criterion, regardless of which criterion was used to select them. The cAIC and

BIC scores of the different schemes were generally similar across criteria (Fig. 2B-C), as expected, with cAIC- and BIC-based schemes achieving the best cAIC and BIC scores, respectively. In contrast, the mAIC scores of mAIC-based schemes were substantially better than those of cAIC- or BIC-based schemes. This difference was particularly apparent across all protein datasets and the green plant plastid DNA datasets.

### mAIC adds modest runtime and memory overhead

We measured the runtime of each PartitionFinder configuration on 50 CPU threads, using the same runs as above (Fig. 3A-B). Because mAIC computation is an additional step relative to cAIC- and BIC-based merging, we assessed its contribution to the total runtime, which depends on the data type. For the DNA datasets, mAIC computation accounted for a notable fraction of the total runtime (8-27%), because substitution model estimation is fast for nucleotide data. For the protein datasets, mAIC computation accounted for only a small fraction of the total runtime (1-2%), as the final model estimation step on protein partitions dominates the overall cost. As a result, cAIC-based PartitionFinder had the longest overall runtime on protein datasets: it retains more partitions and thus must do more computation in the final step, which selects the best model for each partition.

**Fig. 3.**
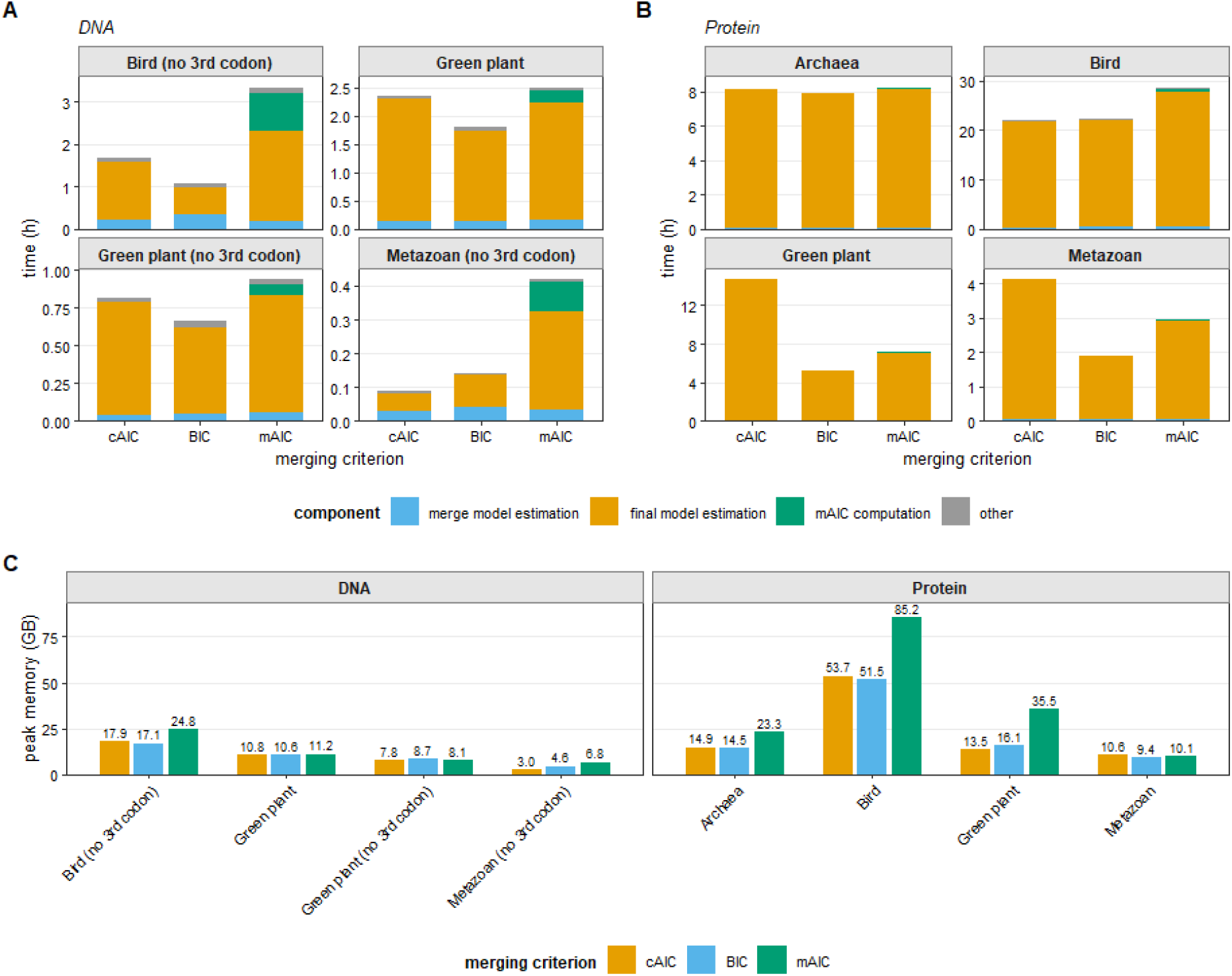
Runtime and memory usage of the PartitionFinder relaxed clustering algorithm under different merging criteria. **A, B**: Runtime breakdown for DNA (**A**) and protein (**B**) datasets. Stacked bars show the time spent on each component of PartitionFinder. **C**: Peak memory usage by the PartitionFinder relaxed clustering algorithm. Numbers above each bar indicate values in GB.

Peak memory usage varied across datasets (Fig 3C). For most datasets, PartitionFinder with mAIC showed a higher memory peak than with cAIC or BIC. However, for some datasets (e.g. green plant DNA, metazoan protein), mAIC did not increase peak memory relative to cAIC and BIC.

### mAIC has a different optimum along the merging path

To examine how each information criterion behaves across the full merging path, defined as the sequence of partitioning schemes obtained by successively merging partition pairs, we ran the PartitionFinder greedy algorithm on a 40-locus subset of the metazoan DNA dataset, merging partitions sequentially from 40 down to 1 (Fig. 4).

**Fig. 4.**
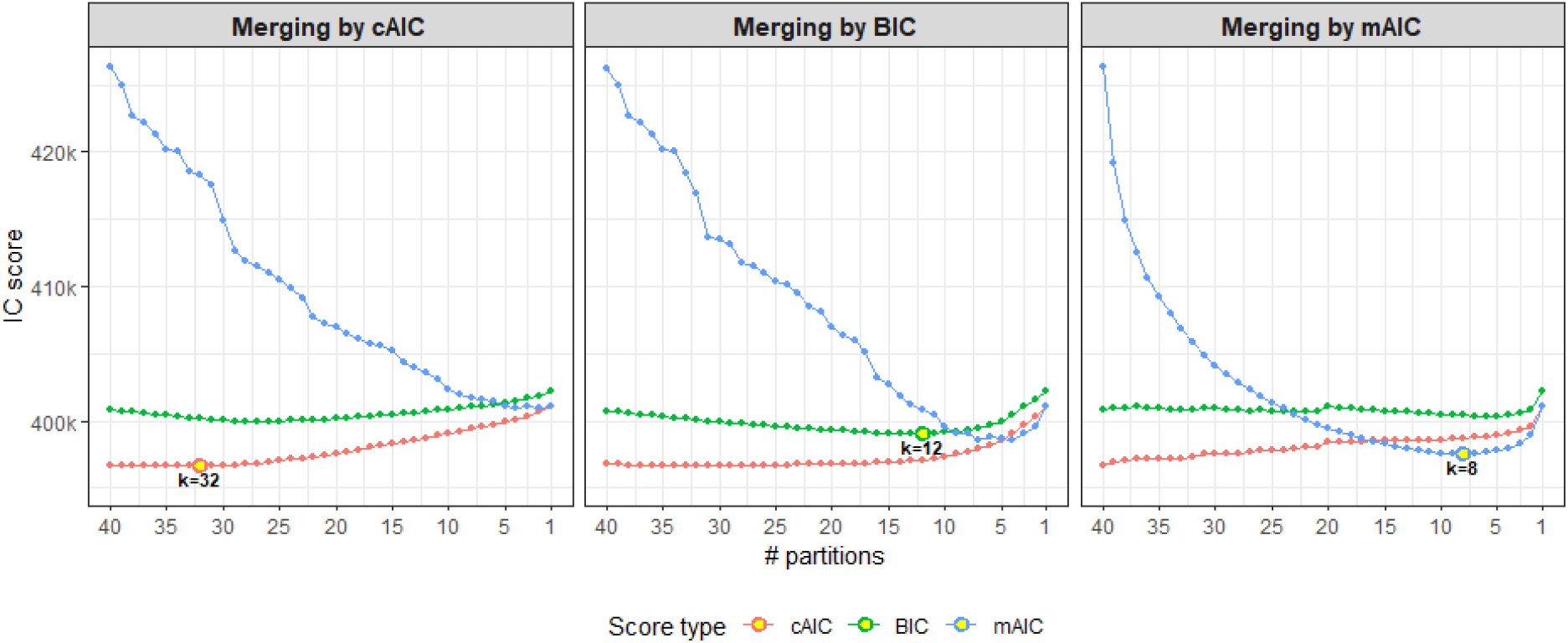
Information criterion scores along the greedy merging path on a 40-locus metazoan DNA subset, from 40 partitions down to 1. Each panel corresponds to a different merging criterion. Colours indicate the IC score type. The highlighted point in each panel marks the optimal score and the corresponding number of partitions (k) for that panel’s merging criterion.

When mAIC was used as the merging criterion, the mAIC score decreased markedly as partitions were merged and reached a clear minimum at 8 partitions. In contrast, when cAIC or BIC was used as the merging criterion, the corresponding scores changed more gradually and reached their optima much earlier along the merging path, at 32 and 12 partitions, respectively. Notably, the mAIC curve under cAIC- (Fig. 4 left panel) or BIC-based merging (Fig. 4 middle panel) did not reach as low a minimum as under mAIC-based merging (Fig. 4 right panel), confirming that optimising the partitioning scheme using the cAIC or BIC does not simultaneously optimise mAIC.

### mAIC-based partitioning schemes influence tree topologies and bootstrap supports of green plant phylogenies

We inferred the green plant phylogenies with 1000 ultrafast bootstrap (UFBoot) replicates (Hoang et al. 2018) under the partitioning schemes estimated by different PartitionFinder criteria (Fig. 2) for the protein, DNA, and DNA excluding third codon positions alignments. For convenience, we refer to the resulting trees as the cAIC tree, BIC tree, and mAIC tree, and compare their topologies with the published tree of Ruhfel et al. (2014), estimated under a no-merge partitioning scheme. That study highlighted several key nodes across plant evolution where topologies are controversial or bootstrap supports were low. Since Ruhfel et al. (2014) used standard bootstrapping (SBS), whose support values are not directly comparable to UFBoot values (UFBoot and SBS are calibrated differently, with UFBoot support of ∼95% and SBS support of ∼70% respectively taken as thresholds for strong support; Minh et al. 2013), we re-inferred the ML trees under their original partitioning scheme with 1000 UFBoot replicates to enable a fair comparison, hereafter referred to as the no-merge tree.

In total, we have 15 trees, including our 12 trees inferred under 3 datasets and 4 partitioning schemes and 3 published trees. Figure 5, Supplementary Figures S8 and S9 show the tree inferred under the mAIC-based scheme for protein, DNA, and DNA excluding third codon positions, respectively. The trees inferred under other criteria are almost identical to the mAIC trees and available in Data, Code and Software Availability section. All our trees share the same backbone with the published trees, differing only at a few contentious nodes (Ruhfel et al. 2014). In the following we describe these differences to illustrate the effect of different partitioning schemes; we do not attempt to make any biological interpretation about green plant evolution.

**Fig. 5.**
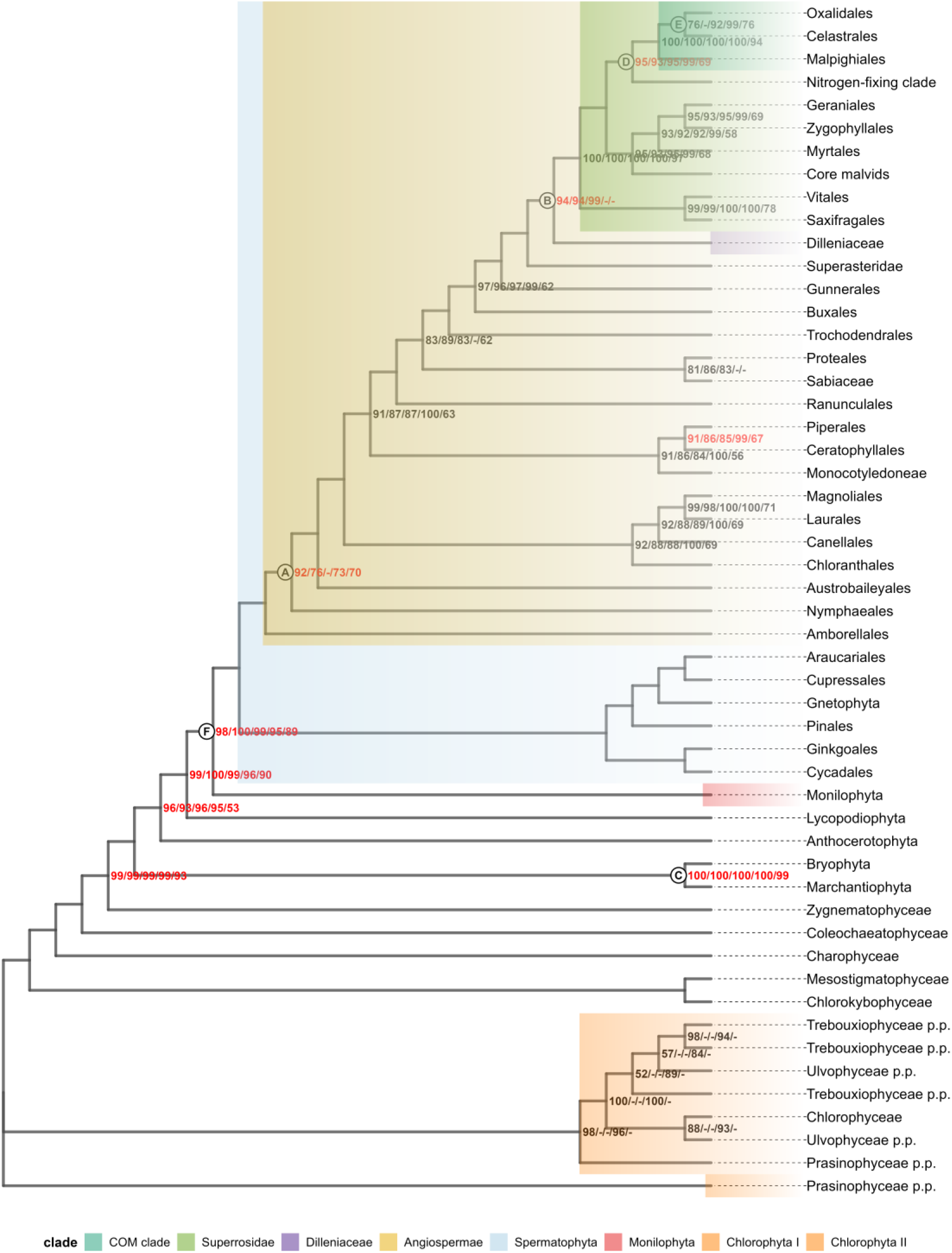
Maximum Likelihood tree inferred from the green plant plastid protein alignment under the mAIC-based partitioning scheme. Bootstrap support values at each node are shown in the format mAIC/BIC/cAIC/no-merge/published tree, where the first four values are ultrafast bootstrap (UFBoot) supports from trees inferred under the corresponding PartitionFinder-mAIC, -BIC, -cAIC and without merging, whereas the fifth value denotes the standard bootstrap (SBS) support from the published tree. A dash (-) indicates that the corresponding clade was not recovered in that tree. Nodes without annotation have 100% supports across all five values, hence not shown. Red numbers highlight key backbone nodes discussed in the original study. Letters A-F mark the nodes presented in the Figure 6. Coloured blocks denote clades where topological disagreements were observed among the trees inferred under different merging criteria or data type in our study.

Bootstrap supports of the contested nodes are highlighted in red (Fig. 5 and Supplementary Figs S8-S9) and further summarised in Figure 6 across all data types. There are two main discrepancies. First, except for the cAIC protein tree, all other trees place *Amborellales* at the base of *Angiospermae* (Fig. 6A, top topology), hence in agreement with the published tree. Support for this *Amborellales*-sister relationship was also higher in the mAIC tree (UFBoot: 92%) than in the BIC (UFBoot: 76%) and no-merge (UFBoot: 73%) trees. Second, except for the no-merge and published protein trees, all other trees placed *Dilleniaceae* as sister to *Superrosidae* (Fig. 6B, top topology). For both placements of *Amborellales* (Fig. 6A) and *Dilleniaceae* (Fig. 6B), the DNA trees (with or without third codon positions) show consistently higher supports (near 100%) than protein trees.

**Fig. 6.**
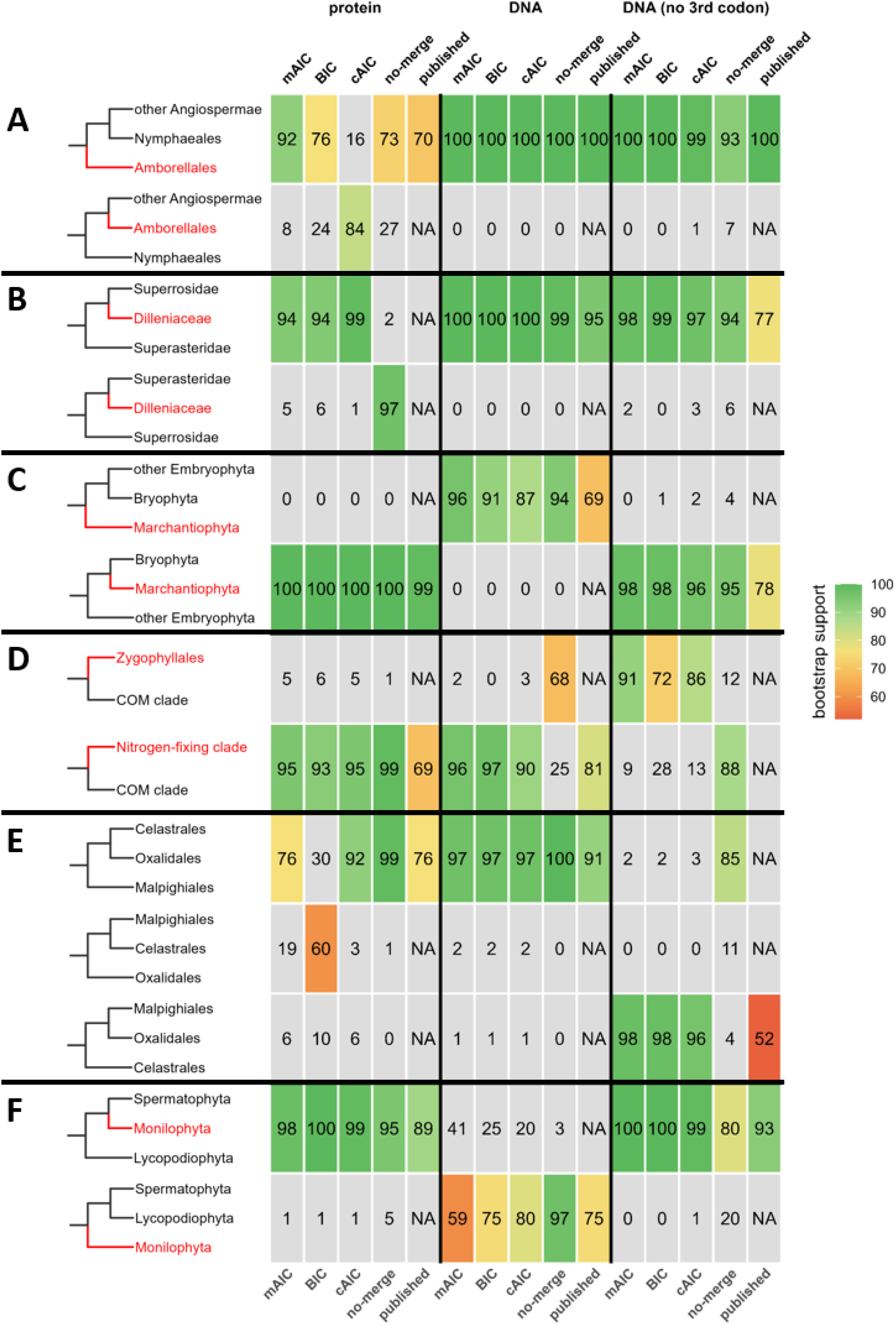
Bootstrap supports for alternative resolutions of each key node of green plant evolution under the different PartitionFinder merging criteria (mAIC, BIC, cAIC), the no-merge partitioning scheme, and the published trees, for each of the three data types. For our own trees, support for every alternative resolution could be recovered from the bootstrap replicates; for Ruhfel et al. (2014), only the bootstrap consensus tree was available, so a standard bootstrap value is shown only for the resolution present in that tree, and no value is given for any alternative where their consensus was unresolved at the node (e.g. the protein trees in **B**). Note that the ultrafast bootstrap (UFBoot; in mAIC/BIC/cAIC/no-merge trees) and standard bootstrap (SBS; in published trees) support values shown in the heatmap are not directly comparable due to different interpretations (Minh et al. 2013).

At these and other key nodes (Fig. 6A-D), the mAIC trees recovered the published relationships with consistently high support. This was the case for the placement of *Marchantiophyta* and *Bryophyta* (Fig. 6C), and for the relationships among the COM clade (*Celastrales*, *Oxalidales* and *Malpighiales*), nitrogen-fixing clade, and *Zygophyllales* (Fig. 6D). Although the topology recovered at these nodes differed between data types, support for the recovered topology remained high in the mAIC trees (UFBoot: 91–98%), whereas it was more variable in the BIC (UFBoot: 72-98%), cAIC (UFBoot: 86-96%) and no-merge (UFBoot: 68-95%) trees.

The mAIC trees received the lower support at some nodes (Fig. 6E-F). For example, within the COM clade (Fig. 6E), *Oxalidales* was recovered as sister to *Celastrales* in the mAIC (UFBoot: 76%), cAIC (92%) and no-merge (99%) protein trees (Fig. 6E, top topology). Whereas the BIC protein tree instead placed *Celastrales* as sister to *Malpighiales* (UFBoot: 60%; Fig. 6E, middle topology). The DNA trees agree with the protein tree with very high supports (UFBoot: 97-100%). Interestingly, the DNA tree excluding third codon positions recovered a third topology, where *Oxalidales* is sister to *Malpighiales* (Fig. 6E, bottom topology), under all three merging criteria (UFBoot: 96-98%). For the next key node (Fig. 6F), *Monilophyta* was placed as sister to *Spermatophyta* with high UFBoot supports in protein and DNA excluding third codon positions like the published tree (Fig. 6F, top topology). Whereas the DNA trees show a monophyletic clade of *Spermatophyta* and *Lycopodiophyta* (Fig. 6F, bottom topology) with low UFBoot supports (mAIC: 59%, BIC: 75%, cAIC: 80%), but still agreeing with the published DNA tree.

Overall, the mAIC trees always corroborate the published trees with higher supports than other information criteria, no matter what data types are used.

## Discussion

In this study, we integrated the mAIC into the PartitionFinder implementation in IQ-TREE 3, extending both the relaxed clustering and greedy search strategies to use mAIC as the merging criterion. We evaluated the new approach on simulated and empirical datasets spanning DNA and protein data.

Across all analyses, mAIC-based merging consistently simplified partitioning schemes with fewer partitions than cAIC and BIC (Fig. 1 and 2). This behaviour is explainable from the mathematical properties of the marginal likelihood. Under conditional likelihood, each site is evaluated only under the model of the partition to which it is assigned. Under marginal likelihood, each site’s likelihood is a weighted average over the models of all partitions. A substitution model fitted to a specific partition may describe the sites in that partition well but fit sites in other partitions poorly, worsening the overall marginal likelihood. Merging two such partitions into one with a more general model can therefore improve the marginal likelihood, even if the conditional likelihood of either partition decreases slightly. This explains why mAIC tends to accept more merges and produce schemes with fewer partitions. BIC yields fewer partitions than cAIC. This is expected, as BIC applies a heavier penalty on the number of parameters, though both criteria are based on conditional likelihoods.

The merging path of mAIC is different from that of cAIC or BIC. As shown in Fig. 4, each criterion reaches its optimum at a different number of partitions. mAIC-based merging produces a sharp optimum in the mAIC score, whereas the mAIC curve under cAIC- or BIC-based merging does not reach as low a minimum, indicating that optimising cAIC or BIC does not simultaneously optimise mAIC. This is further supported by our correlation analysis (Supplementary Fig. S1): no single metric, including ΔcAIC, was a strong proxy for ΔmAIC throughout the merging process. These results confirm that direct computation of mAIC is necessary for optimising partition merging under this criterion.

We extend the PartitionFinder algorithm (Lanfear et al. 2017) as implemented in IQ-TREE (Chernomor et al. 2016) to additionally compute the mAIC score during merging. As described in the Materials and Methods, we preserve the existing filtering and sorting design, as our correlation analysis did not reveal a better metric for filtering or sorting, and add an mAIC evaluation when a pair is to be merged.

In practice, applying mAIC-based PartitionFinder to the empirical datasets tested in this study (Table 1) only increased the total runtime by a maximum of 27% of extra mAIC computations (Fig. 3). This is because the final model estimation step of PartitionFinder in IQ-TREE runs a full ModelFinder search (Kalyaanamoorthy et al. 2017) on each partition, evaluating all available substitution matrices and free-rate models (Yang 1995; Soubrier et al. 2012), which is particularly expensive. This also explains why cAIC-based PartitionFinder sometimes showed the longest total runtime on protein datasets, as it retained the most partitions and thus required the most model estimation.

According to Susko et al. (2026), mAIC is expected to provide a partitioning scheme that improves the global model parameters, such as species tree topology and branch lengths. In our simulations, however, the improvement in tree inference under mAIC was limited (Supplementary Figs. S2-S7). On the empirical green plant data, by contrast, where inference focuses on specific contested nodes, the mAIC-based partitioning scheme did influence both tree topology and branch support (Figs. 5 and 6; Supplementary Figs. S8 and S9).

The green plant plastid dataset of Ruhfel et al. (2014) is a useful test case for our method: several backbone nodes remain contentious despite large-scale phylogenomic efforts (One Thousand Plant Transcriptomes Initiative 2019). At two of these nodes, independent evidence indicates which resolution is currently well-accepted, so the criteria can be judged against an external standard. The first is the placement of *Amborellales* (Fig. 6A). *Amborellales* alone is widely accepted as the basal *Angiospermae* (Fig. 6A, top topology). Whereas the alternative hypotheses of basal *Nymphaeales* (Fig. 6A, bottom topology) and an *Amborellales* + *Nymphaeales* clade (Xi et al. 2014) have been attributed to limited taxon-sampling and long-branch attraction (Drew et al. 2014). In the protein trees (Fig. 5), mAIC supported the accepted *Amborellales*-first resolution most strongly of the four schemes (UFBoot 92%), whereas cAIC instead recovered the *Nymphaeales*-first alternative (UFBoot 84%). The second is the placement of *Monilophyta* (Fig. 6F). *Monilophyta* + *Spermatophyta* (Fig. 6F, top topology) is strongly supported by nuclear and morphological data (Pryer et al. 2001; Wickett et al. 2014), whereas the conflicting *Lycopodiophyta* + *Spermatophyta* grouping (Fig. 6F, bottom topology) is attributed to compositional bias in plastid genomes (Cox et al. 2014). All four schemes recovered this artefactual topology from the DNA data (Supplementary Fig. S8), but support for it was lowest under mAIC (UFBoot 59%) and near-maximal under the no-merge partitioning scheme (UFBoot 97%). This artefactual topology was absent from every protein (Fig. 5) and DNA excluding third codon positions (Supplementary Fig. S9) tree, consistent with a nucleotide-level artefact. At both nodes, mAIC gave the strongest support to the accepted resolution and the weakest support to the artefactual one.

The remaining focal nodes (Fig. 6B-E), all identified as contentious or weakly supported by (Ruhfel et al. 2014), have no accepted resolution, so no criterion can be judged correct there. They nonetheless show the same sensitivity: both the recovered topology and its support varied with the merging criterion and with the data type, mAIC giving the strongest support at some nodes and the weakest at others, and at several nodes incompatible resolutions each received high support in different analyses.

Our aim here is not to study green plant evolution, and we draw no biological conclusions from these differences. We want to emphasise that data types, models and information criteria can each change the recovered topology, sometimes with high supports for conflicting resolutions. The robustness of a given node cannot be judged from a single analysis. We recommend that practitioners examine the evolution of their species of interests under all settings.

In conclusion, we have efficiently implemented mAIC as a partition-merging criterion in the PartitionFinder algorithm of IQ-TREE 3. On both simulated and empirical data, mAIC consistently selected simpler partitioning schemes than the existing criteria, potentially reducing overparameterisation. At the same time, mAIC matches or improves topological accuracy. We recommend including mAIC in partitioned phylogenomic analyses, as a complement to the existing criteria.

## Data, Code and Software Availability

PartitionFinder-mAIC is available in IQ-TREE version 3.1.4 at https://iqtree.github.io. The analyses in this study were run with modified IQ-TREE builds as described in the Materials and Methods. These programs, together with the datasets, scripts and IQ-TREE output files used in this study, are available at https://doi.org/10.6084/m9.figshare.33433609.

## Supplementary Materials

Supplementary material is available at [supplementary file link]

