## Supplementary for "PartitionFinder-mAIC: Phylogenetic Partitioning using Marginal Akaike Information Criterion"

### Supplementary Materials for "PartitionFinder-mAIC: Phylogenetic Partitioning using Marginal Akaike Information Criterion"

Huaiyan Ren[1,*], Thomas K.F. Wong[1,2], Changsen Jiang[3], Edward Susko[4], Robert Lanfear[3,$], Bui Quang Minh[1,$]

[1] *School of Computing, College of Systems & Society, Australian National University, Canberra, ACT 2600, Australia.*

[2] *Mathematical Sciences Institute, College of Systems & Society, Australian National University, Canberra, ACT 2600, Australia.*

[3] *Ecology and Evolution, Research School of Biology, College of Science & Medicine, Australian National University, Canberra, ACT 2600, Australia.*

[4] *Department of Mathematics and Statistics, Faculty of Science, Dalhousie University, NS, Canada.*

[$] **Co-senior author**

#### Supplementary Figures

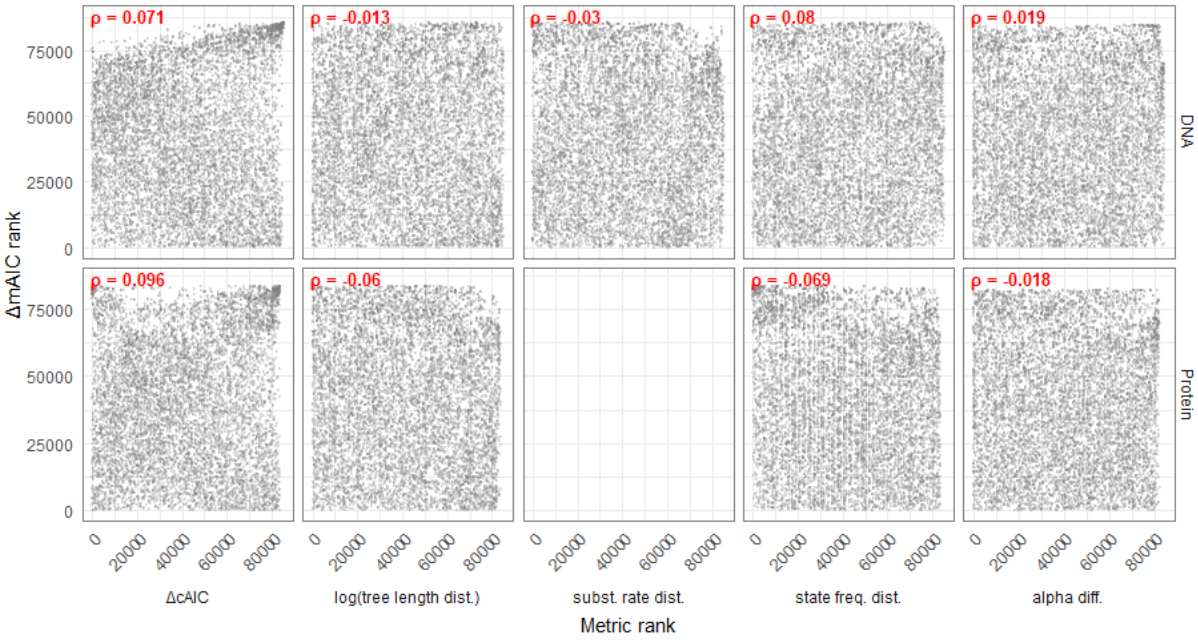


**Fig. S1** Rank–rank scatter plots of ΔmAIC against each tested metric across greedy merging iterations, pooled across data subsets and datasets. Rows correspond to DNA (top) and protein (bottom) data. Columns correspond to the five tested metrics. Each panel displays 8000 randomly sampled data points. Spearman's ρ is shown in each panel.

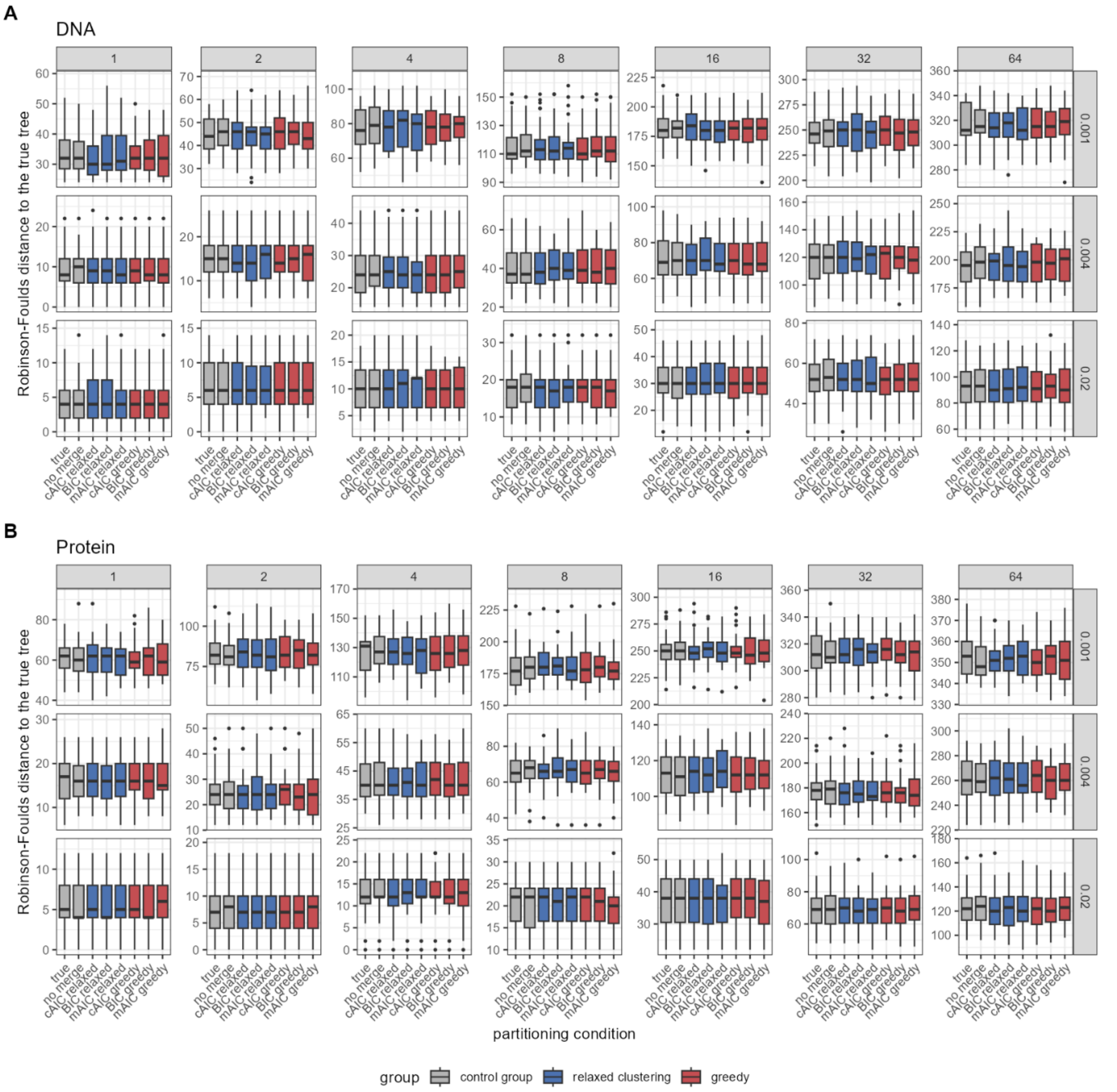


**Fig. S2** Robinson-Foulds distance to the true tree by each criterion and partition-merging strategy under simulation. (**A**) DNA and (**B**) protein datasets. Within each panel, boxplots show the RF distance across replicates, faceted by mean branch length (rows) and external-to-internal branch-length ratio (columns). Asterisks denote significant differences from paired Wilcoxon signed-rank tests, with each criterion compared against mAIC (blue, relaxed clustering; red, greedy). P-values were adjusted for multiple comparisons using the Holm method ($*$ $P < 0.05$, $**$ $P < 0.01$, $***$ $P < 0.001$).

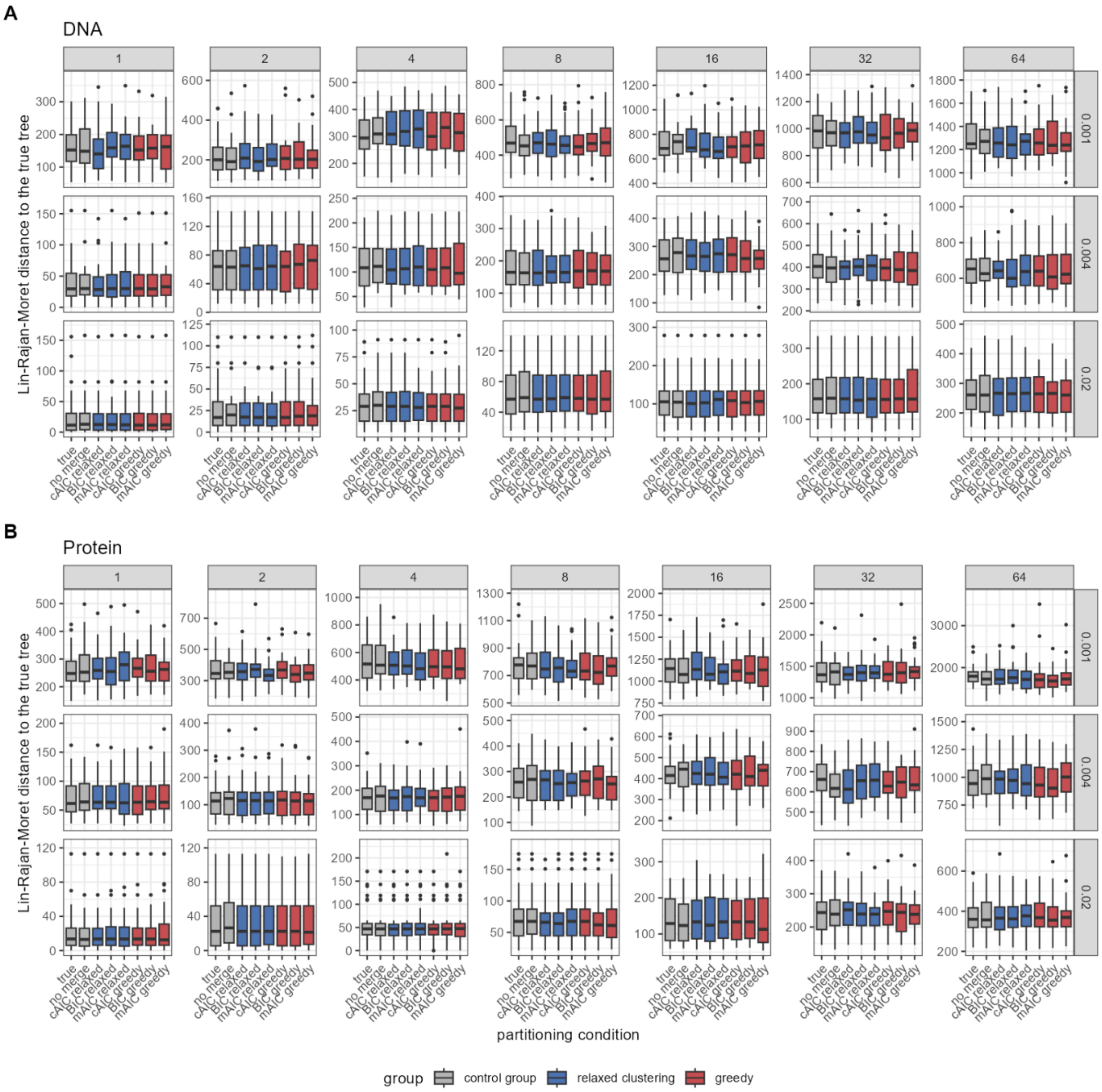


**Fig. S3** Lin-Rajan-Moret distance to the true tree by each criterion and partition-merging strategy under simulation. (**A**) DNA and (**B**) protein datasets. Within each panel, boxplots show the LRM distance across replicates, faceted by mean branch length (rows) and external-to-internal branch-length ratio (columns). Asterisks denote significant differences from paired Wilcoxon signed-rank tests, with each criterion compared against mAIC (blue, relaxed clustering; red, greedy). P-values were adjusted for multiple comparisons using the Holm method (∗ $P < 0.05$, ∗∗ $P < 0.01$, ∗∗∗ $P < 0.001$).

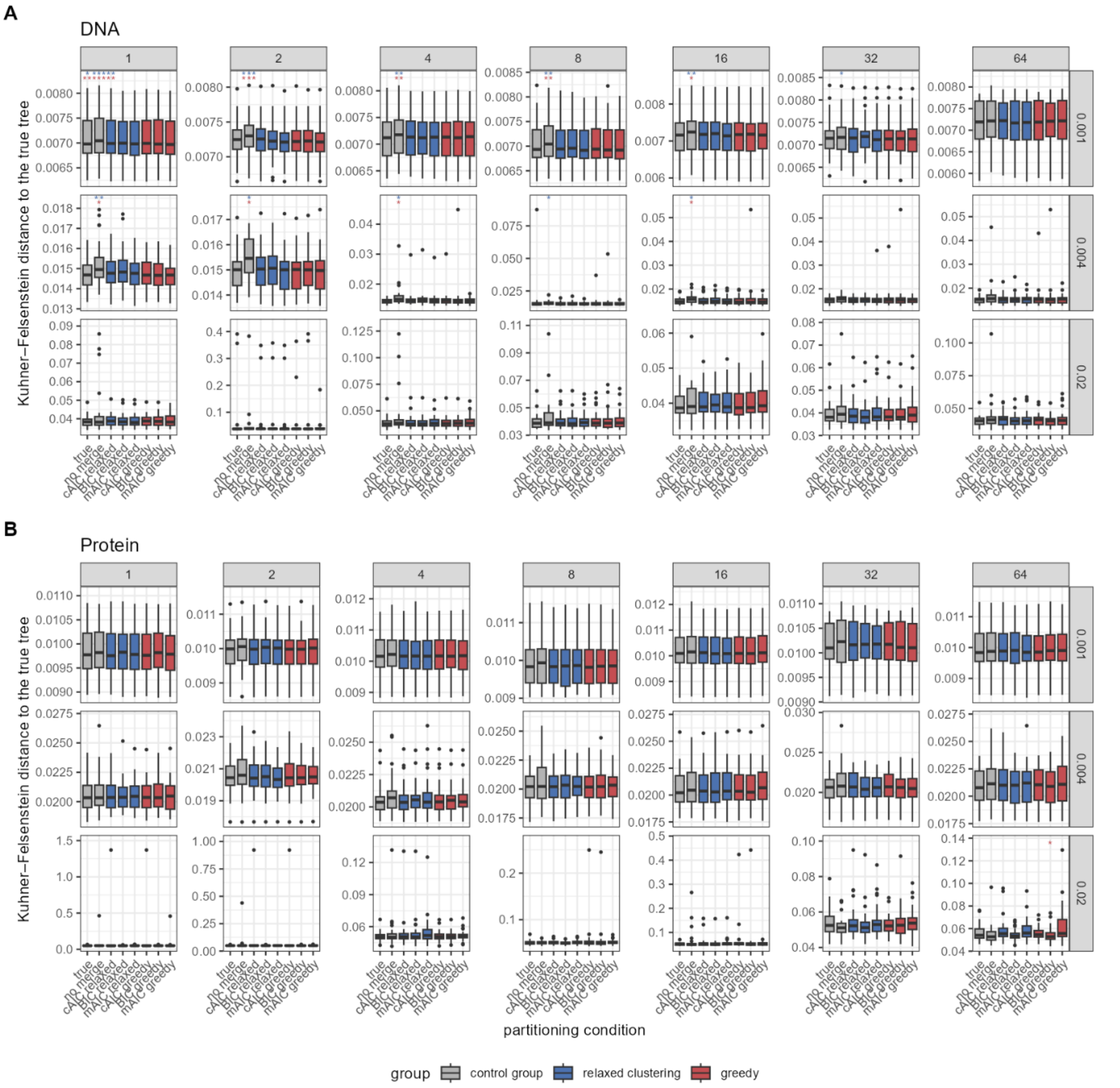


**Fig. S4** Kuhner-Felsenstein distance to the true tree by each criterion and partition-merging strategy under simulation. (**A**) DNA and (**B**) protein datasets. Within each panel, boxplots show the KF distance across replicates, faceted by mean branch length (rows) and external-to-internal branch-length ratio (columns). Asterisks denote significant differences from paired Wilcoxon signed-rank tests, with each criterion compared against mAIC (blue, relaxed clustering; red, greedy). P-values were adjusted for multiple comparisons using the Holm method ($*$ P < 0.05, $**$ P < 0.01, $***$ P < 0.001).

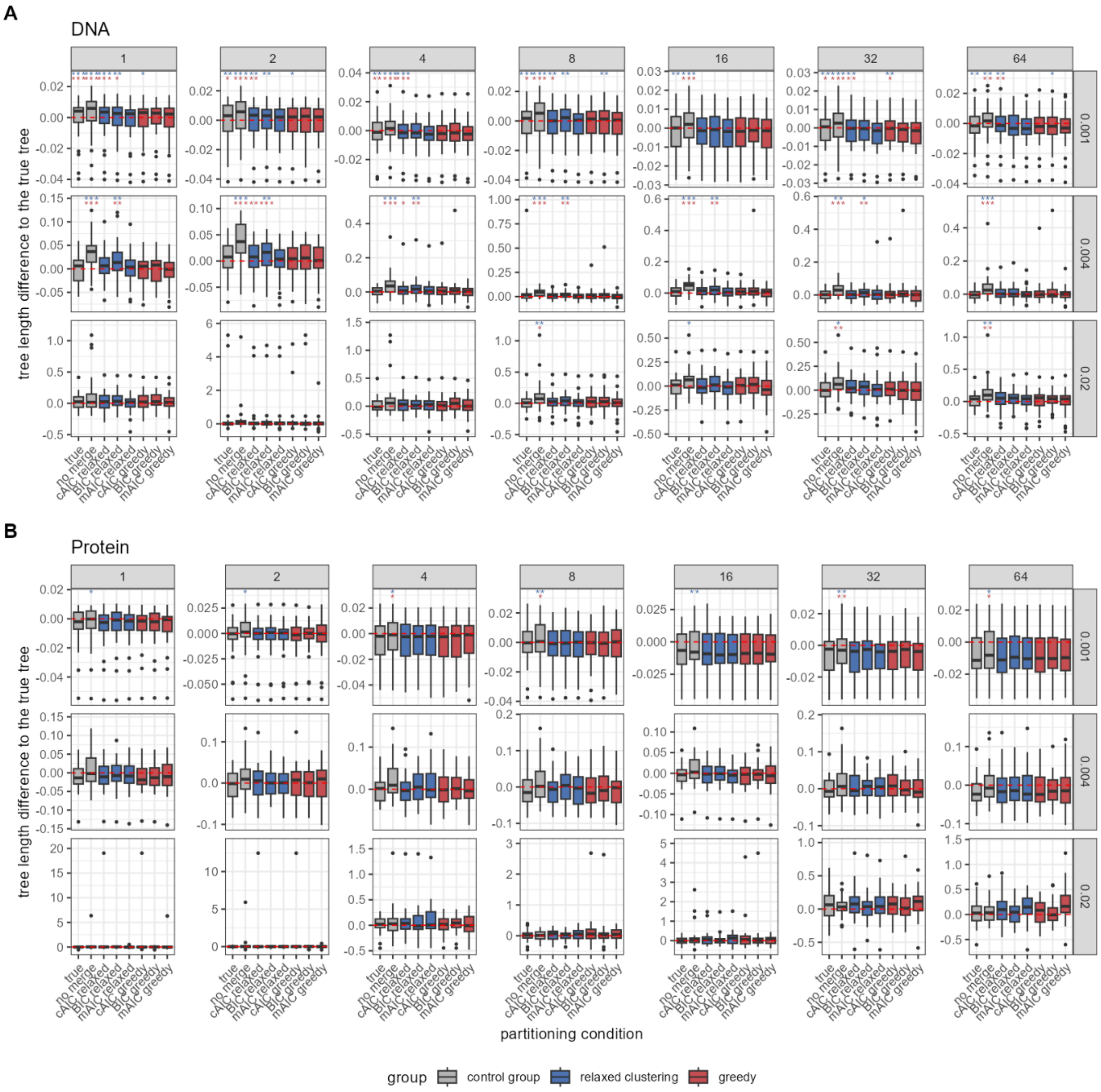


**Fig. S5** Tree length difference to the true tree by each criterion and partition-merging strategy under simulation. (**A**) DNA and (**B**) protein datasets. Within each panel, boxplots show the difference in total tree length from the true tree across replicates, faceted by mean branch length (rows) and external-to-internal branch-length ratio (columns). Red dashed lines indicate the difference is zero. Asterisks denote significant differences from paired Wilcoxon signed-rank tests, with each criterion compared against mAIC (blue, relaxed clustering; red, greedy). P-values were adjusted for multiple comparisons using the Holm method ($*$ $P < 0.05$, $**$ $P < 0.01$, $***$ $P < 0.001$).

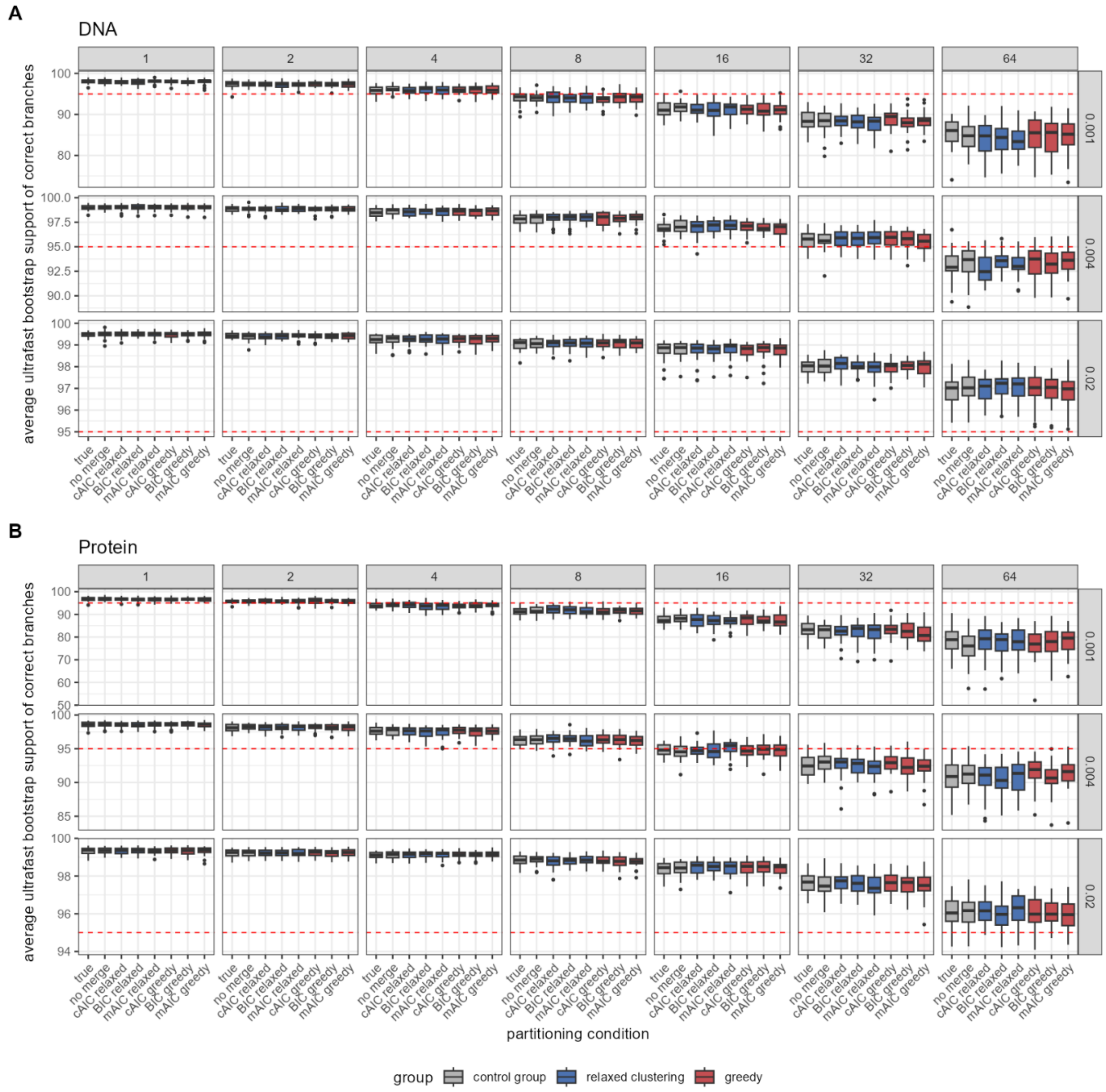


**Fig. S6** Average ultrafast bootstrap support of correct branches by each criterion and partition-merging strategy under simulation. (**A**) DNA and (**B**) protein datasets. Within each panel, boxplots show the average UFBoot support of correctly inferred branches across replicates, faceted by mean branch length (rows) and external-to-internal branch-length ratio (columns). Red dashed lines indicate the bootstrap support is 95. Asterisks denote significant differences from paired Wilcoxon signed-rank tests, with each criterion compared against mAIC (blue, relaxed clustering; red, greedy). P-values were adjusted for multiple comparisons using the Holm method ($*$ $P < 0.05$, $**$ $P < 0.01$, $***$ $P < 0.001$).

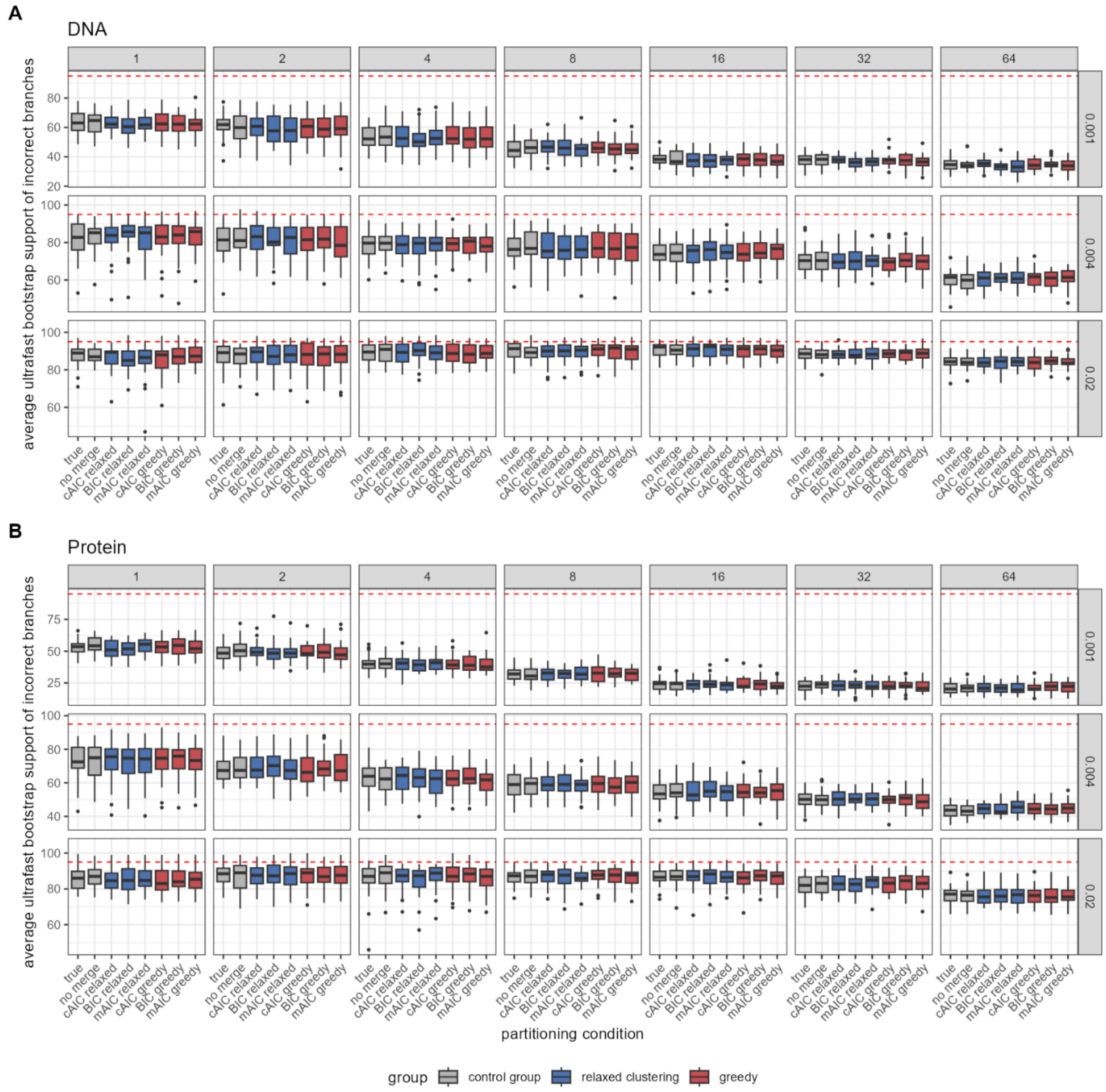


**Fig. S7** Average ultrafast bootstrap support of incorrect branches by each criterion and partition-merging strategy under simulation. (**A**) DNA and (**B**) protein datasets. Within each panel, boxplots show the average UFBoot support of incorrectly inferred branches across replicates, faceted by mean branch length (rows) and external-to-internal branch-length ratio (columns). Red dashed lines indicate the bootstrap support is 95. Asterisks denote significant differences from paired Wilcoxon signed-rank tests, with each criterion compared against mAIC (blue, relaxed clustering; red, greedy). P-values were adjusted for multiple comparisons using the Holm method ($*$ $P < 0.05$, $**$ $P < 0.01$, $***$ $P < 0.001$).

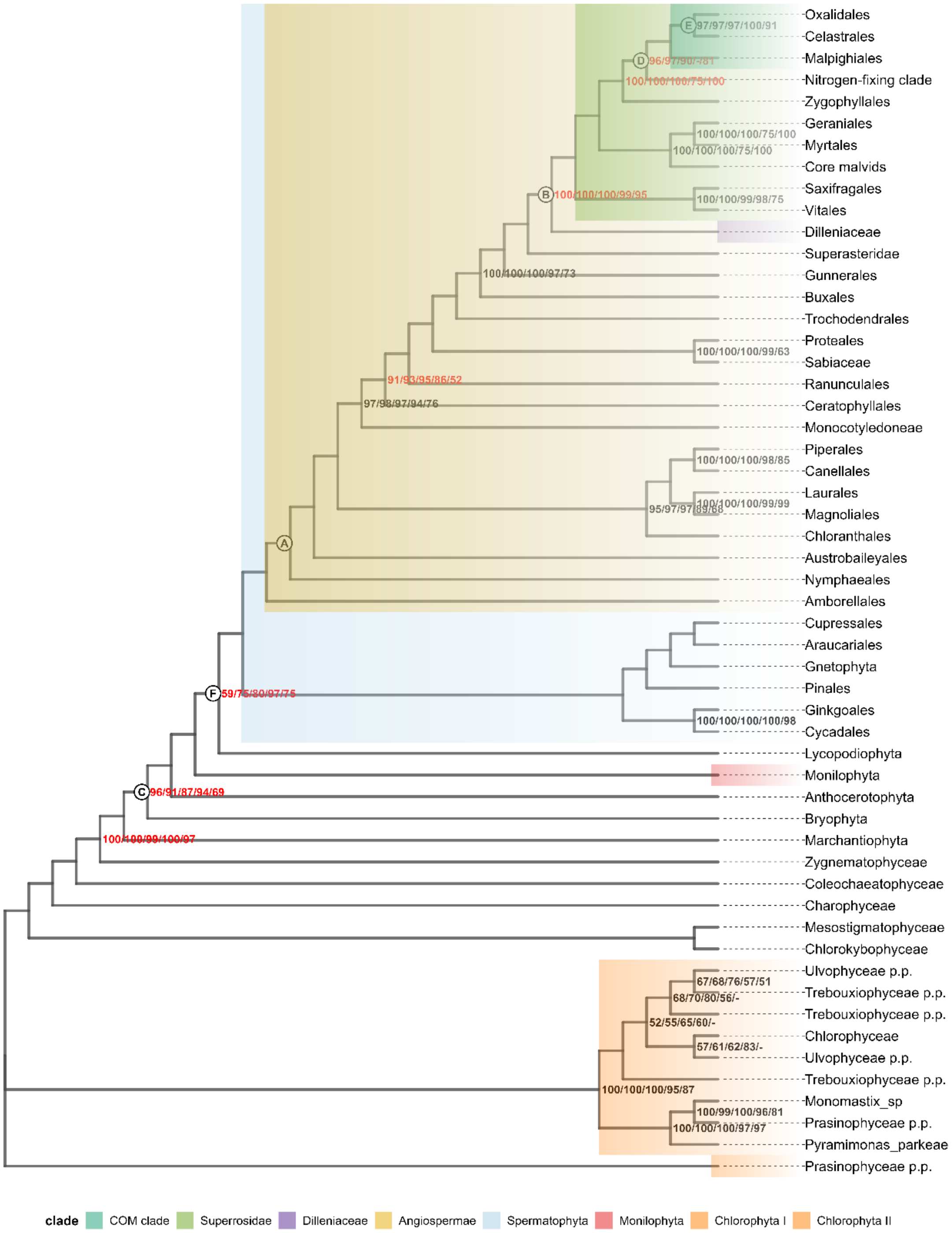


**Fig. S8** Maximum Likelihood tree inferred from the green plant plastid DNA alignment with all codon positions under the mAIC-based partitioning scheme. Bootstrap support values at each node are shown in the format mAIC/BIC/cAIC/no-merge/published tree, where the first four values are ultrafast bootstrap (UFBoot) supports from trees inferred under the corresponding PartitionFinder-mAIC, -BIC, -cAIC and without merging, whereas the fifth value denotes the standard bootstrap (SBS) support from the published tree. A dash (-) indicates that the corresponding clade was not recovered in that tree. Nodes without annotation have 100% supports across all five values, hence not shown. Red numbers highlight key backbone nodes discussed in the original study. Letters A-F mark the nodes presented in the Figure 6. Coloured blocks denote clades where topological disagreements were observed among the trees inferred under different merging criteria or data type in our study.

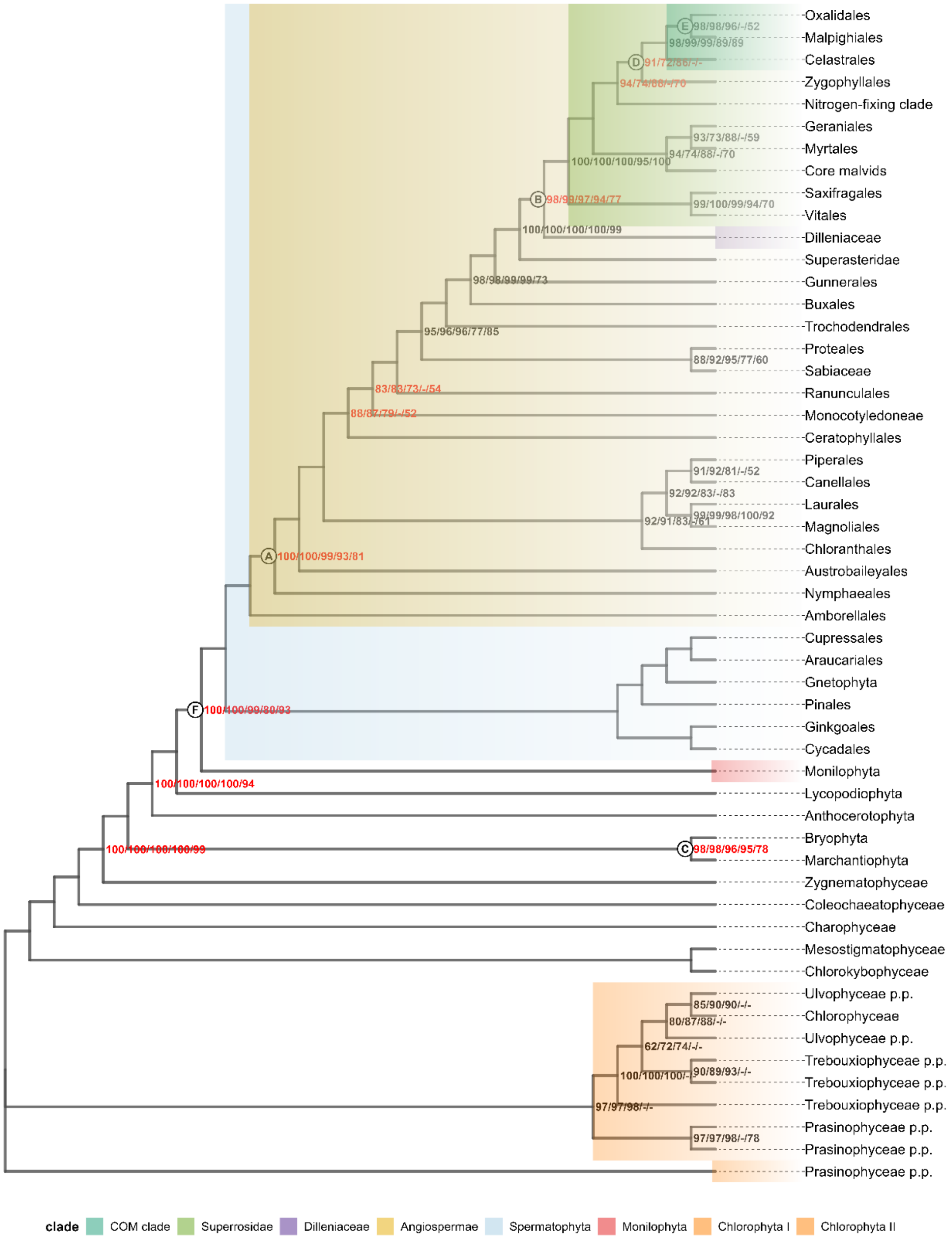


**Fig. S9** Maximum Likelihood tree inferred from the green plant plastid DNA alignment excluding third codon positions under the mAIC-based partitioning scheme. Bootstrap support values at each node are shown in the format mAIC/BIC/cAIC/no-merge/published tree, where the first four values are ultrafast bootstrap (UFBoot) supports from trees inferred under the corresponding PartitionFinder-mAIC, -BIC, -cAIC and without merging, whereas the fifth value denotes the standard bootstrap (SBS) support from the published tree. A dash (-) indicates that the corresponding clade was not recovered in that tree. Nodes without annotation have 100% supports across all five values, hence not shown. Red numbers highlight key backbone nodes discussed in the original study. Letters A-F mark the nodes presented in the Figure 6. Coloured blocks denote clades where topological disagreements were observed among the trees inferred under different merging criteria or data type in our study.
